# Astrocytic Chromatin Remodeler ATRX Gates Hippocampal Memory Consolidation through Metabolic and Synaptic Regulation

**DOI:** 10.1101/2025.07.21.665940

**Authors:** Miguel A. Pena-Ortiz, Julia K. Sunstrum, Alireza Ghahramani, Haley McConkey, Vanessa Dumeaux, Wataru Inoue, Nathalie G. Bérubé

## Abstract

Astrocytes are increasingly recognized as active regulators of synaptic transmission and memory, yet the epigenetic mechanisms underlying their contribution to cognitive processes remain poorly defined. Here, we investigated the role of the chromatin remodeler ATRX in astrocytes by generating mice with inducible, astrocyte-specific *Atrx* deletion (aiKO) using tamoxifen administration at postnatal days 10–12, resulting in ATRX loss in approximately half of hippocampal and cortical astrocytes. Transcriptomic profiling of hippocampal tissue at one and three months revealed a progressive increase in differentially expressed genes, with early enrichment for cytoskeletal and immune pathways and later dysregulation of energy metabolism, ion transport, and synaptic gene sets. Electrophysiological recordings from CA1 pyramidal neurons in aiKO slices demonstrated increased neuronal excitability, reduced membrane capacitance, and decreased frequency of spontaneous excitatory postsynaptic currents, indicating non-cell-autonomous neuronal dysfunction. Morphological analysis identified a transient reduction in dendritic branching at one month and a selective loss of thin dendritic spines by three months, without changes in total dendrite length or overall spine density. Behaviorally, aiKO mice displayed normal locomotion, anxiety, and short-term memory, but exhibited deficits in 24-hour novel object recognition and long-term spatial memory in the Morris water maze. These findings demonstrate that ATRX-mediated chromatin remodeling in astrocytes is essential for maintaining hippocampal transcriptional homeostasis, neuronal function, and long-term memory. Our results highlight a critical role for astrocytic epigenetic regulation in cognitive processes and suggest that astrocyte dysfunction may contribute to the pathogenesis of ATR-X syndrome and related intellectual disability disorders, underscoring the importance of targeting multiple cell types for therapeutic intervention.

**Highlights:**

- Astrocytic ATRX loss causes non-cell-autonomous neuronal hyperexcitability
- Inducing ATRX deficiency in astrocytes causes selective long-term memory deficits
- Dynamic transcriptomic changes reveal metabolic and synaptic pathway dysregulation

## Introduction

Astrocytes, the most abundant glial cell type in the mammalian brain, have emerged as active regulators of synaptic transmission, circuit plasticity, and memory formation, extending beyond their conventional roles in structural support and metabolic maintenance[1]. Their functional diversity is shaped by complex developmental programs and region-specific cues, with epigenetic mechanisms now recognized as central to determining astrocyte identity, maturation, and specialized functions[2, 3]. Recent advances have highlighted the importance of chromatin remodeling and epigenetic regulation in astrocyte heterogeneity and their capacity to influence neuronal circuits, yet the molecular mechanisms by which these processes in astrocytes contribute to cognitive function remain poorly understood, representing a significant gap in our understanding of memory and intellectual disability disorders.

ATR-X syndrome, caused by mutations in the *ATRX* gene, is characterized by moderate to severe intellectual disability, seizures, white matter abnormalities, and diverse developmental defects [4, 5]. ATRX encodes an ATP-dependent chromatin remodeling factor that binds to specific histone marks and partners with the DAXX histone chaperone to facilitate the incorporation of the histone variant H3.3 at telomeres and pericentric heterochromatin [6]. Beyond its canonical roles in maintaining constitutive heterochromatin and genome integrity, ATRX modulates gene expression through mechanisms such as preventing G-quadruplex formation during transcriptional elongation, promoting long-range chromatin interactions, and recruiting Polycomb complexes to neurodevelopmental targets [7–10]. Despite these established functions, the role of ATRX in developing and mature astrocytes, and the consequences of its loss for neuronal physiology and cognitive abilities, remain largely unexplored. This is particularly relevant as astrocyte dysfunction is increasingly implicated in neurodevelopmental disorders, and astrocytes are now appreciated as critical regulators of memory processes.

Given the recognized influence of astrocytes on neurodevelopment and cognitive function, we sought to determine how ATRX dysfunction in astrocytes affects neuronal function and memory *in vivo*. To address this, we generated and characterized mice with inducible, conditional inactivation of ATRX in astrocytes during the early postnatal period, a developmental window that allows us to investigate both the maturation and maintenance roles of ATRX. Our approach enabled us to assess the non-cell-autonomous effects of ATRX-null astrocytes on hippocampal CA1 neuron physiology and to link these changes to specific deficits in long-term spatial and recognition memory. Transcriptomic profiling revealed dynamic, progressive changes in metabolic, ionic, and synaptic pathways in the hippocampus, implicating astrocytic ATRX in the maintenance of transcriptional homeostasis required for neuronal function. These findings provide new insight into the epigenetic regulation of astrocyte function, establish a mechanistic link between glial chromatin remodeling and cognitive processes, and implicate astrocytic ATRX as a critical modulator of neural circuits underlying memory.

## Results

### Selective deletion of ATRX in astrocytes does not disrupt astrocyte density or morphology

To generate mice with astrocyte-specific deletion of ATRX, we utilized an inducible Cre-loxP system (Slc1a3-cre/ERT;AtrxloxP or Glast-creERT;AtrxloxP [11, 12]) and administered tamoxifen daily from postnatal day 10 to 12, creating what we refer to as ATRX <u>a</u>strocyte-inducible knockout (ATRX aiKO) mice (**Fig. 1A**). The efficiency of ATRX recombination in astrocytes was assessed by immunofluorescence staining for ATRX and the astrocyte marker GFAP at one month of age. Quantitative analysis showed that approximately 50% of GFAP^+^ astrocytes in the hippocampus lack ATRX expression in aiKO mice (**Fig. 1B,C**). Since GFAP does not label all astrocytes, particularly in the cortex, we also stained for S100β, which confirmed that 40-50% of astrocytes in both the cortex and hippocampus are ATRX-negative in the aiKO mouse brain (**Fig. 1D,E**). To verify the specificity of Cre-mediated recombination, we performed co-staining for ATRX and the oligodendrocyte marker OLIG2 or the microglia marker IBA1. The majority of OLIG2-positive and IBA1-positive cells retained ATRX expression in the cortex and hippocampus of ATRX aiKO mice (**Fig. S1A-D**). Notably, astrocyte density, as assessed by S100β staining, was unchanged between ATRX aiKO and control mice in both cortex and hippocampus (**Fig. 1F**).

**Figure 1:**
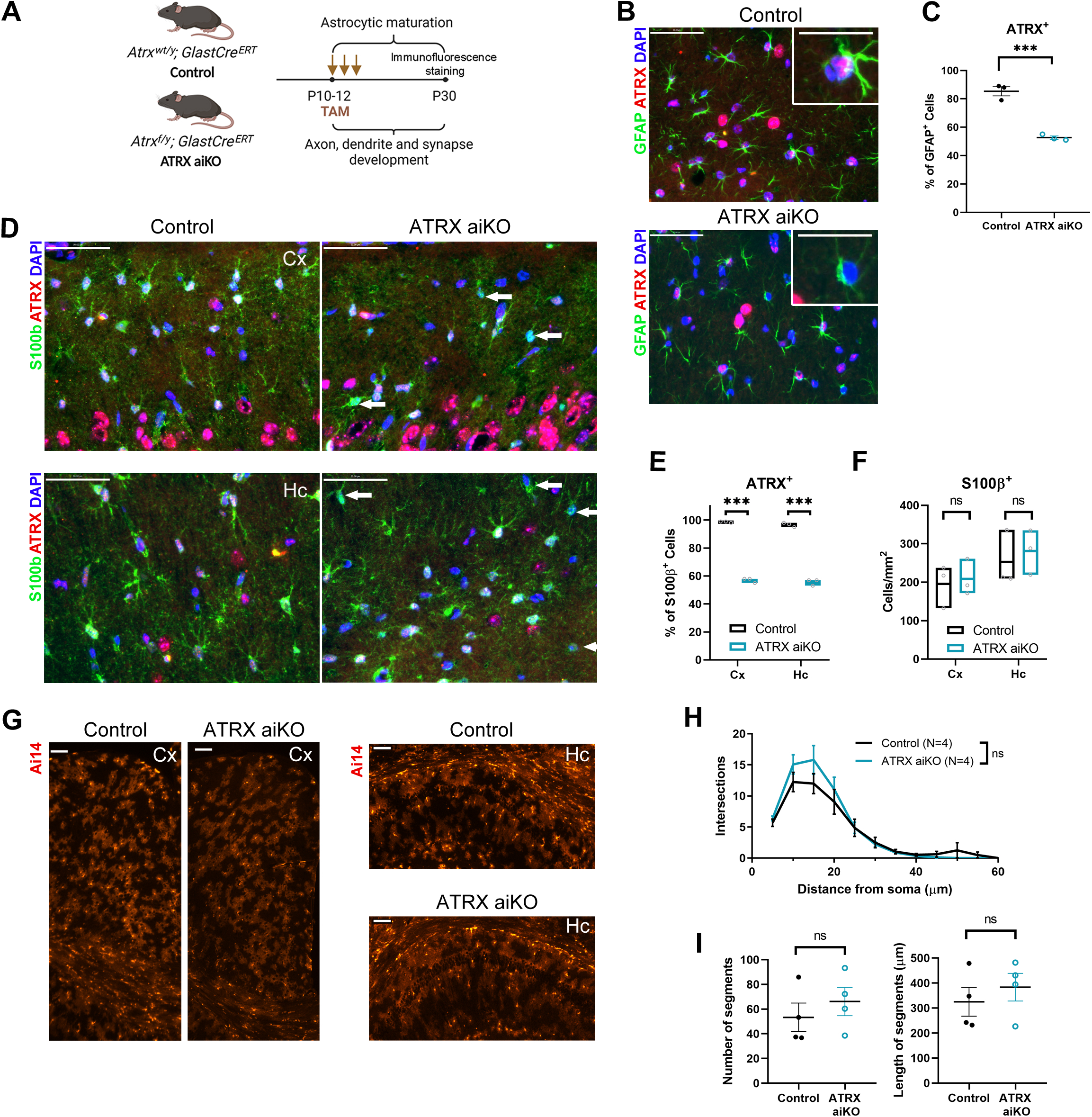
Efficiency of Cre-mediated ATRX deletion in astrocytes upon tamoxifen treatment. (A) Control (Atrxwt/y;GlastCreERT) and ATRX aiKO (Atrxf/y;GlastCreERT) mice were injected with tamoxifen i.p. daily from P10 to P12 and brain sections analyzed by immunofluoresence staining at P30. (B) Immunofluorescence staining of ATRX and the astrocyte marker GFAP in the hippocampus of ATRX aiKO and control mice. (C) Quantification of ATRX+/GFAP+ cells in the hippocampus. Unpaired Student’s t-test, t_(4)_=9.610. (D) Immunofluorescence staining of brain cryosections for ATRX and the astrocyte marker S100β in the cortex and hippocampus. Arrows point to ATRX-null cells. (E) Quantification of ATRX+ /S100β+ astrocytes in the cortex and hippocampus. Unpaired Student’s t-test, t_Cx(4)_=9.610, t_Hc(4)_=26.65. (F) Quantification of S100β+ cells in the cortex and hippocampus reveals similar astrocyte density in control and ATRX aiKO mice. Unpaired Student’s t-test, t_Cx(4)_=0.2929, t_Hc(4)_=0.5276. (G) Ai14 fluorescence of ATRX aiKO and control Cre+ astrocytes in cortical and hippocampal sections. (H) Sholl analysis of GFAP Staining of Ai14+ astrocytes at P90, N=4 mice from each genotype. twANOVA, FGenotype (1,72)=1.559, p=0.2159. (I) Dendrite number and length of GFAP projections at P90. N=4 mice each genotype. Number of segments, t(6)=0.7876, p=0.4609; length of segments t(6)=0.7325, p=0.4915. Scalebars, (B,D) 50 µm; (G), 100 µm. Cx: Cortex, Hc: Hippocampus. Data shown as +/- SEM, unpaired Student’s t-test, ***p<0.001, n=3 animals per genotype.

To further evaluate astrocyte morphology and coverage, we utilized the Ai14 Cre reporter allele to fluorescently label Cre-expressing astrocytes and found that the overall distribution and coverage of tdTomato-positive astrocytes was similar in both genotypes (**Fig. 1G**). For a more quantitative assessment, we performed Sholl analysis on GFAP-stained, Ai14-positive astrocytes using confocal microscopy. Although there was a trend toward increased branching close to the soma (at approximately 10 μm), there was no significant difference in overall branching complexity between control and ATRX-null astrocytes (**Fig. 1H**). Likewise, the number and length of GFAP-positive processes were not significantly different between groups (**Fig. 1I**).

Taken together, these results demonstrate that inducible deletion of ATRX in astrocytes using postnatal tamoxifen administration achieves efficient recombination in approximately half of the astrocyte population in the cortex and hippocampus, with high specificity for astrocytes over other glial cell types. Importantly, loss of ATRX in astrocytes does not result in major morphological changes, suggesting that ATRX is dispensable for the maintenance of basic astrocyte structure under baseline conditions in the postnatal brain.

### Progressive and dynamic transcriptomic dysregulation in the hippocampus following astrocytic ATRX deletion

To investigate the impact of ATRX deletion in astrocytes on hippocampal gene expression, we performed bulk RNA sequencing on hippocampal tissue from ATRX aiKO and control mice at one and three months of age.

At one month, we identified 928 differentially expressed genes (DEGs; from transcripts with aggregated Lancaster p value <0.05; see Methods), with nearly equal proportions of upregulated (49.25%) and downregulated (50.75%) genes. Most of these changes exhibit moderate fold changes, as visualized by volcano plot (**Fig. 2A**). Gene ontology analysis of DEGs at this early timepoint revealed significant enrichment in biological processes related to cytoskeletal regulation, including “negative regulation of microtubule depolymerization” and “microtubule bundle formation” (**Fig. 2B**). Additional enriched categories include inflammatory and immune processes such as “positive regulation of toll-like receptor 3 signaling pathway” and “positive regulation of NF-κB transcription factor activity” as well as processes relevant to calcium signaling and neuronal function, including “positive regulation of cytosolic calcium ion concentration,” “positive regulation of synapse assembly,” “positive regulation of neuron migration,” and “positive regulation of dendritic spine morphogenesis” (**Table S1**).

**Figure 2.**
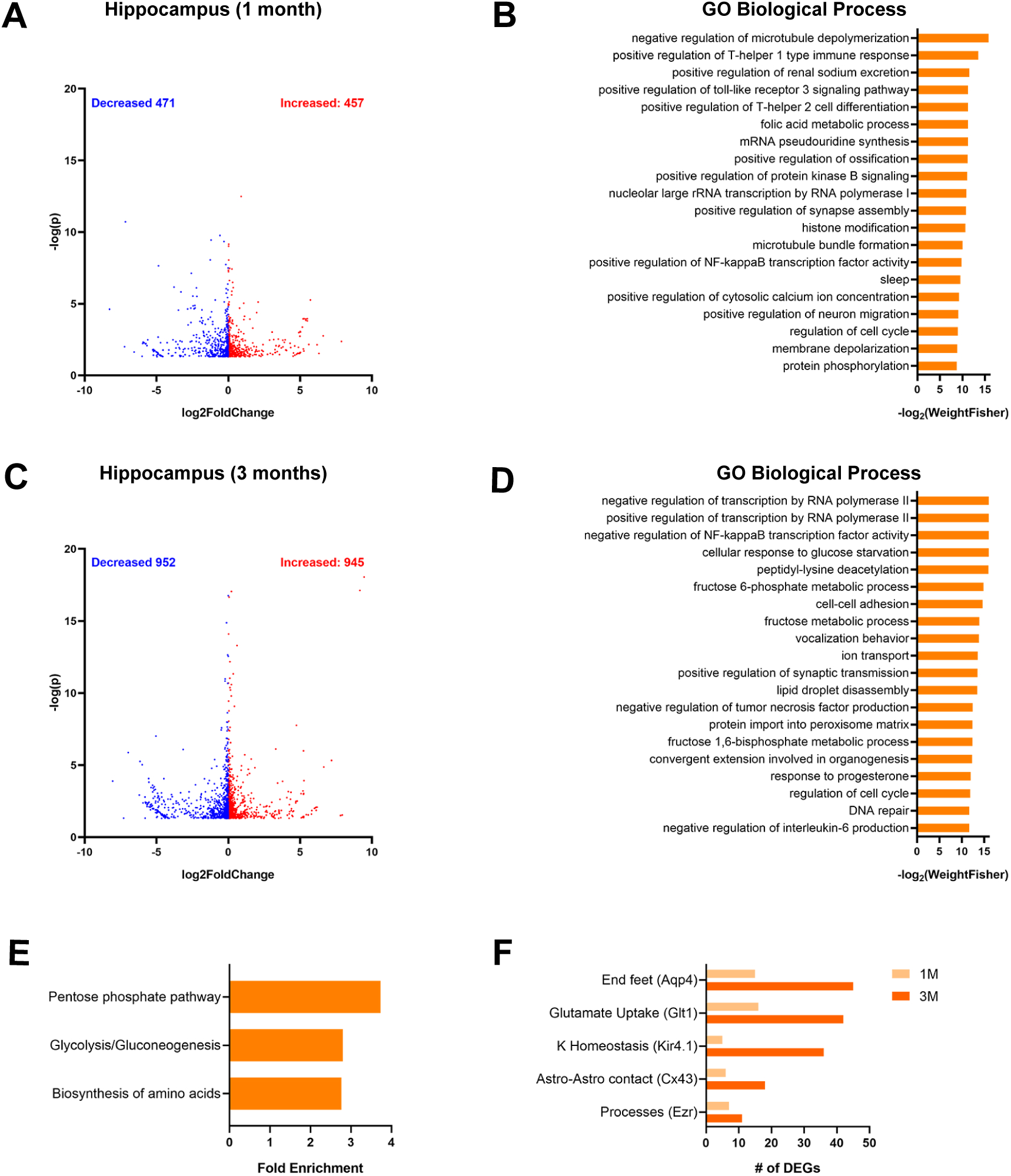
Transcriptomic changes in hippocampal tissue and astrocytes of ATRX aiKO mice. (A) Volcano plot of DEGs and (B) Top 20 most significantly enriched Biological Process Gene Ontology categories at P30. (C) Volcano plot of DEGs and (D) Top 20 most significantly enriched Biological Process Gene Ontology categories at P90. (E) KEGG pathway enrichment analysis of DEG at P90. (F) Integration of our DEGs at P30 and P90 with proteins detected in astrocytic compartments.

At three months, the extent of transcriptional dysregulation increases, with 1,897 DEGs identified (aggregated Lancaster p value <0.05), again with a balanced distribution of increased (49.82%) and decreased (50.18%) gene expression (**Fig. 2C**). Most DEGs at this stage also showed moderate fold changes. Gene ontology analysis at three months revealed a shift in the affected biological processes, with the most highly enriched categories related to transcriptional regulation by RNA polymerase II, as well as several metabolic processes, including “cellular response to glucose starvation,” “fructose 6-phosphate metabolic process,” and “fructose metabolic process” (**Fig. 2D**, **Table S2**). KEGG pathway analysis further supported the enrichment of energy metabolism pathways, such as the pentose phosphate pathway, glycolysis/gluconeogenesis, and amino acid biosynthesis (**Fig. 2E**).

To assess whether these transcriptomic changes include genes relevant to astrocyte-specific functions, we compared our hippocampal DEGs to proteins previously identified in distinct astrocytic compartments using a BioID proximity labeling approach [13]. We found DEGs encoding proteins located in all compartments at one and three months of age with a greater number of deregulated genes at the later timepoint (**Fig 2F**). Notably, the most abundant DEGs at three months correspond to proteins localized to astrocytic endfeet, and over 35 DEGs were associated with proteins in proximity to the glutamate transporter Glt1 and the potassium channel Kir4.1, suggesting potential alterations in glutamate uptake and potassium homeostasis.

In summary, the results suggest that ATRX is required for the maintenance of transcriptional homeostasis in astrocytes and that its loss may contribute to metabolic and synaptic dysfunction in the hippocampus.

### Non-cell-autonomous structural remodeling of CA1 neurons following astrocytic ATRX deletion

To investigate whether ATRX-deficient astrocytes influence neuronal structure, we performed detailed morphological analyses of CA1 pyramidal neurons in the hippocampus of ATRX aiKO and control mice using the Thy1-GFP-M reporter to sparsely label neurons. GFP-positive CA1 neurons were imaged at both one and three months of age to capture potential developmental and persistent effects (**Fig. 3A**)

**Figure 3.**
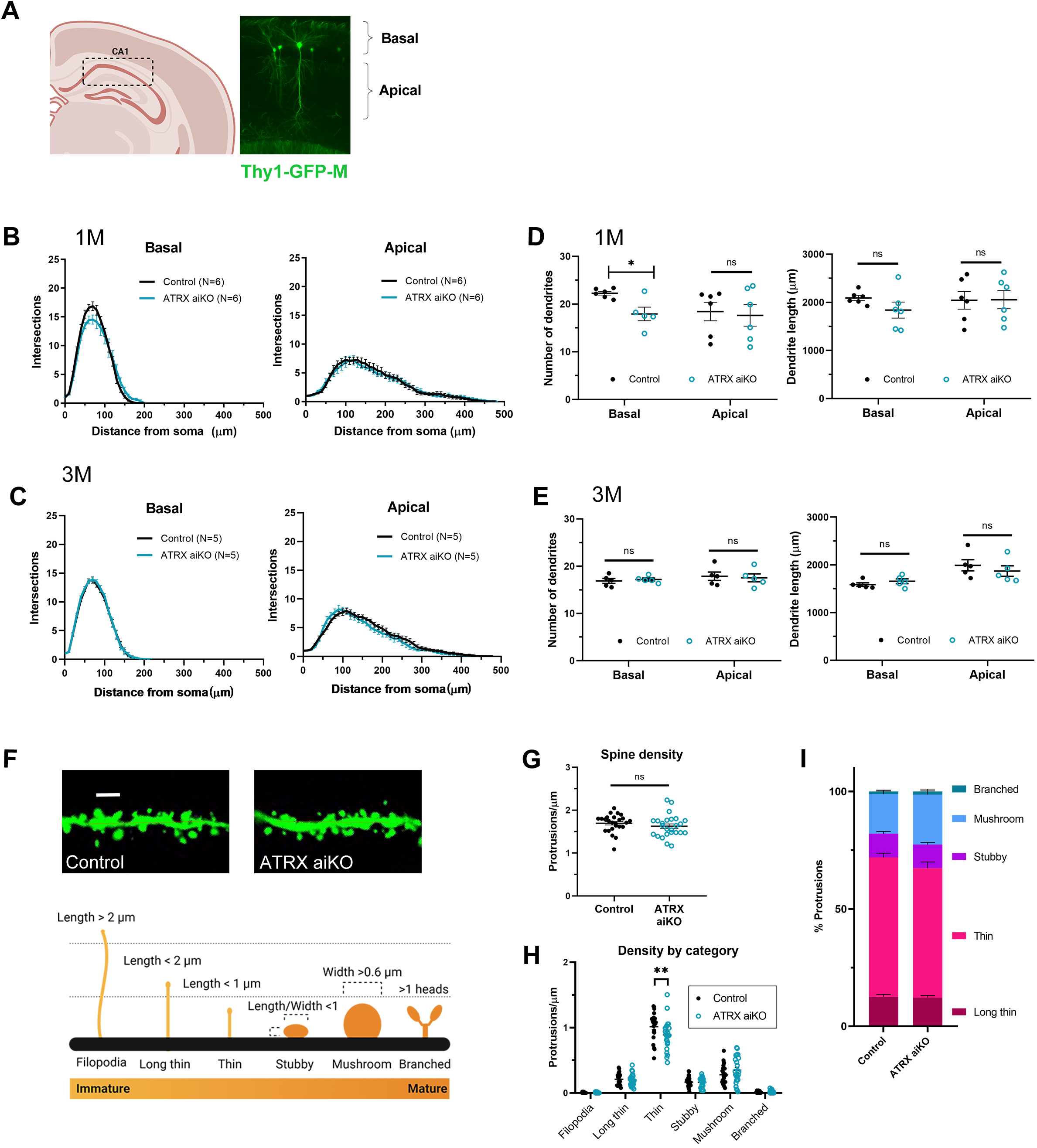
Morphological assessment of CA1 hippocampal neurons in control and ATRX aiKO mice. (A) Representative image of Thy1-GFP+ hippocampal CA1 neuron. (B) Sholl analysis of basal and apical dendrites of CA1 neurons at P30. N=6 mice from each genotype. twANOVA, basal dendrites: F_Genotype (1,10)_=0.4217, p=0.5307; apical dendrites: F_Genotype (1,10)_=0.0285, p=0.869. (C) Sholl analysis of basal and apical dendrites of CA1 neurons at P90. N=5 mice each genotype. twANOVA, basal dendrites: F_Genotype (1,9)_=0.234, p=0.640; apical dendrites: F_Genotype (1,9)_=0.508, p=0.496. (D) Dendrite number and length of CA1 neurons at P30. N=6 mice each genotype. Basal number, t_(9)_=3.219, *p<0.05; apical number, t_(10)_=0.272, p=0.792, basal length, t_(10)_=1.447, p=0.179; apical length, t_(10)_=1.804, p=0.101. (E) Dendrite number and length of CA1 neurons at P90. N=5 mice of each genotype. Basal number, t_(8)_=0.473, p=0.649, apical number, t_(8)_=0.260, p=0.801, basal length, t_(8)_=1.135, p=0.289; apical length, t_(8)_=0.7499, p=0.4748. (F) Representative images of dendritic segments from P90 control and ATRX aiKO hippocampal CA1 neurons. Scale bar, 2um. Diagram of criteria from Risher et al. 2014 used to categorize spines. (G) Average spine density of hippocampal CA1 apical dendrites, t_(48)_=1.017, p=0.314. (H) Spine density by morphology, twANOVA, F_Genotype (1,288)_=0.608, p=0.436; mc, **p<0.01. (I) Proportion of dendritic spine categories of control and ATRX aiKO hippocampal CA1 neurons. twANOVA, F_Genotype (1,288)_=2.579×10^-7^, p=0.9996; multiple comparison: n.s. Average of 5 segments from 5-10 neurons from N=3 mice of each genotype were analysed.

Sholl analysis did not identify differences in dendritic branching between genotypes (**Fig. 3B,C** and **Fig. S2A,B**). However, decreased branching in the range of 50 to 100 µm and in the average maximum number of intersections of CA1 basal dendrites were detected (**Fig. 3B and S2C**). We also identified a significant decrease in the total number of basal dendrites at this early time point (**Fig. 3D**). By three months of age, these structural deficits were no longer observed: the number and length of primary, secondary, and tertiary dendrites were comparable between ATRX aiKO and control mice, and overall dendritic complexity had normalized (**Fig. 3C, 3E, S2B, S2D**). Soma area also remained unchanged between groups at both time points (**Fig. S3E**).

Given that astrocytes can regulate neuronal spines by pruning during development [14–16], spine density and morphology was scored and categorized as long thin, thin, stubby, mushroom or branched [17] (**Fig. 3F**). While the average length and width of dendritic spines were similar between genotypes (**Fig. S2F**), a selective reduction in the density of thin spines was detected in ATRX aiKO mice at three months (**Fig. 3G, 3H**). Total spine density was not significantly altered, but there was a trend toward a decreased proportion of thin spines and a corresponding increase in mushroom spines in the ATRX aiKO group (**Fig. 3I**).

Together, these results demonstrate that loss of ATRX in astrocytes leads to a transient reduction in basal dendritic complexity of CA1 neurons during early postnatal development, followed by a selective and persistent decrease in the density of thin dendritic spines in adulthood.

### Astrocytic ATRX deficiency induces hyperexcitability and reduced synaptic input in CA1 pyramidal neurons

To address the consequences of astrocytic ATRX deletion on neuronal function, we used *ex vivo* patch clamp electrophysiology in brain slices obtained from 3 to 6-month-old ATRX aiKO and control mice (**Fig. 4A**). The results show a pronounced hyperexcitability of CA1 neurons of the ATRX aiKO compared to control hippocampal slices, identified by a significant leftward shift in the firing-current injection relationship (**Fig. 4B**). The hyperexcitability is not accompanied by changes in the action potential amplitude (**Fig. 4C**) or threshold (**Fig. 4D**), suggesting that active membrane properties underlying the action potentials (e.g. voltage-dependent Na^+^, Ca^2+^ and K^+^ channels) remain generally intact in ATRX aiKO. On the other hand, the hyperexcitability was accompanied by a significant decrease in the capacitance of the neurons (**Fig. 4E**), indicating that they are electronically more compact and consequently hyperexcitable [18, 19]. Indeed, capacitance and input resistance (membrane resistance at rest) are negatively correlated (p=0.0082, **Fig. 4F**). However, input resistance alone is not significantly different between aiKO and control neurons (**Fig. 4G**), suggesting additional mechanisms contributing to the subthreshold membrane properties. In contrast to the hyperexcitability of intrinsic properties, we observed decreased excitatory synaptic inputs to CA1 neurons in ATRX aiKO, with a decrease in the frequency but not the amplitude of spontaneous excitatory postsynaptic currents (sEPSCs) (**Fig. 4H-J**). Overall, these results show that CA1 neurons in the vicinity of ATRX-null astrocytes are dysfunctional, exhibiting decreased frequency of excitatory synaptic input but augmented excitability upon stimulation.

**Figure 4.**
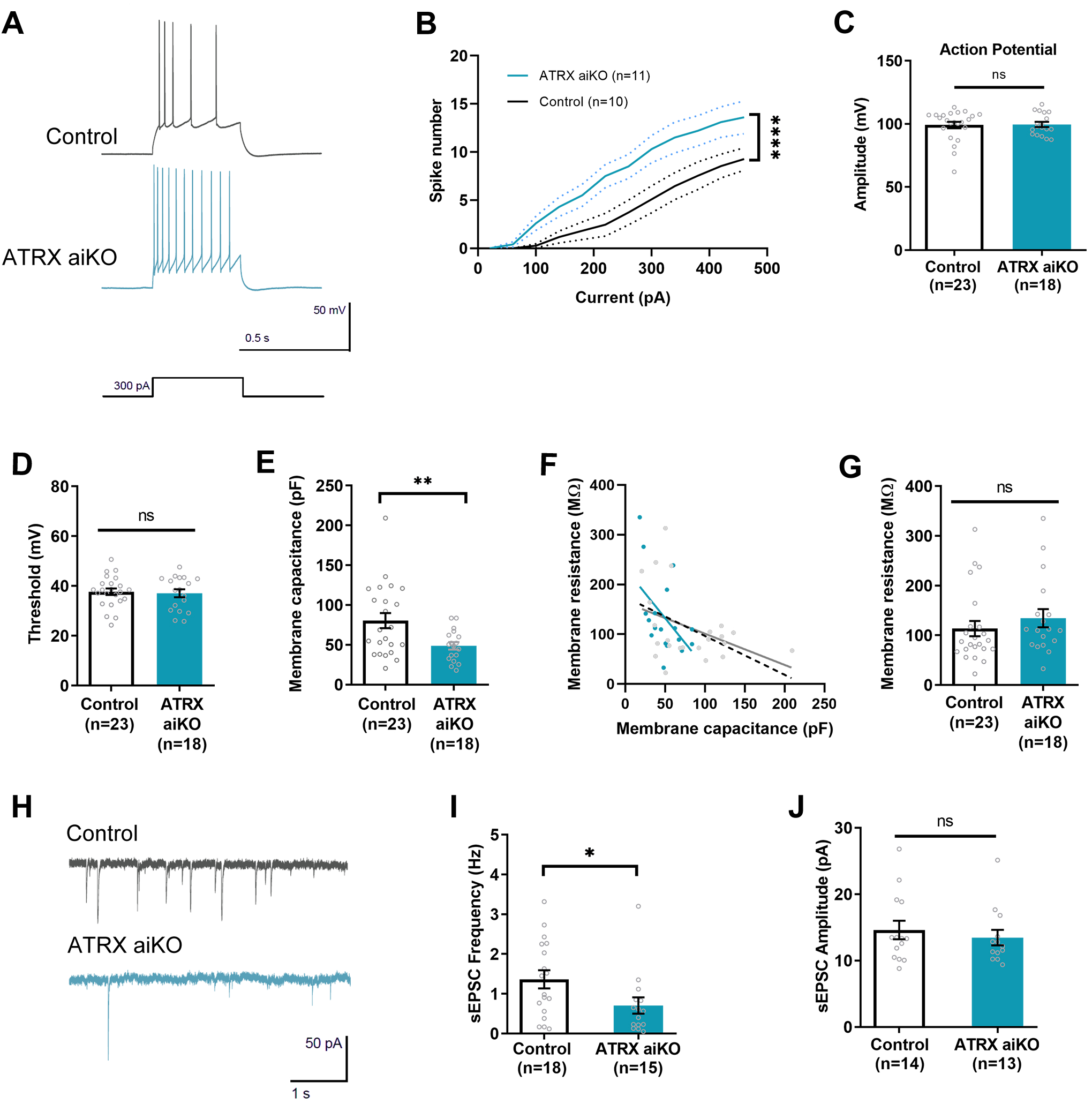
Decreased synaptic function and hyperexcitability in ATRX aiKO hippocampal CA1 neurons. A) Representative action potential firing during 300pA current step in control and ATRX aiKO hippocampal CA1 neurons. (B) Action potential firing as spike number during current step, twANOVA, FGenotype (1,228)=59.21, p<0.0001. (C) Amplitude of rheobase action potential, unpaired Student’s t-test, t(39)=0.1257, p=0.9006. (D) Threshold of rheobase action potential of hippocampal CA1 neurons, unpaired Student’s t-test, t(39)=0.2988, p=0.7667. (E) Membrane capacitance of hippocampal CA1 neurons, unpaired Student’s t-test, t(39)=2.746, **p<0.01. (F) Membrane resistance vs capacitance, linear regression F(1,39)=7.768, p= 0.0082. (G) Membrane resistance of hippocampal CA1 neurons, unpaired Student’s t-test, t(39)=0.8874, p=0.3803. (H) Representative sEPSC traces from hippocampal CA1 neurons in brain sections of control and ATRX aiKO mice. (I) Frequency of sEPSCs, unpaired Student’s t test, t(31)=2.116, *p<0.05. (J) Amplitude of sEPSCs, unpaired Student’s t test, t(25)=0.6281, p=0.5357. Data represented as mean +/- SEM, n=5 mice of each genotype.

### Astrocytic ATRX deficiency selectively impairs long-term spatial and recognition memory

Given the electrophysiological and structural changes observed in CA1 neurons of ATRX aiKO mice, we performed a battery of behavior tests on ATRX aiKO and control mice to evaluate locomotion, anxiety, and various types of memory **(Fig. 5A)**. The results show that ATRX aiKO mice have normal locomotion and activity levels based on the distance travelled in the open field test (**Fig. 5B**). Vertical counts or rearing, an anxiety and exploration indicator in the open field test, as well as time at the center of the arena are also unaltered in ATRX aiKO mice (**Fig. 5B and Fig. S3A**). Similarly, no difference in the time spent in the dark compartment of the light dark box test is detected (**Fig. S3B**). Working memory is also intact based on the comparable number of entries or alternations in the Y-maze task (**Fig. 5C**, **S3C**). Testing contextual fear memory also revealed no differences between control and ATRX aiKO mice (**Fig. 5D**), despite ATRX aiKO mice showing a reduced response in the freezing time immediately after the shock in the training session (**Fig. S3D**).

**Figure 5.**
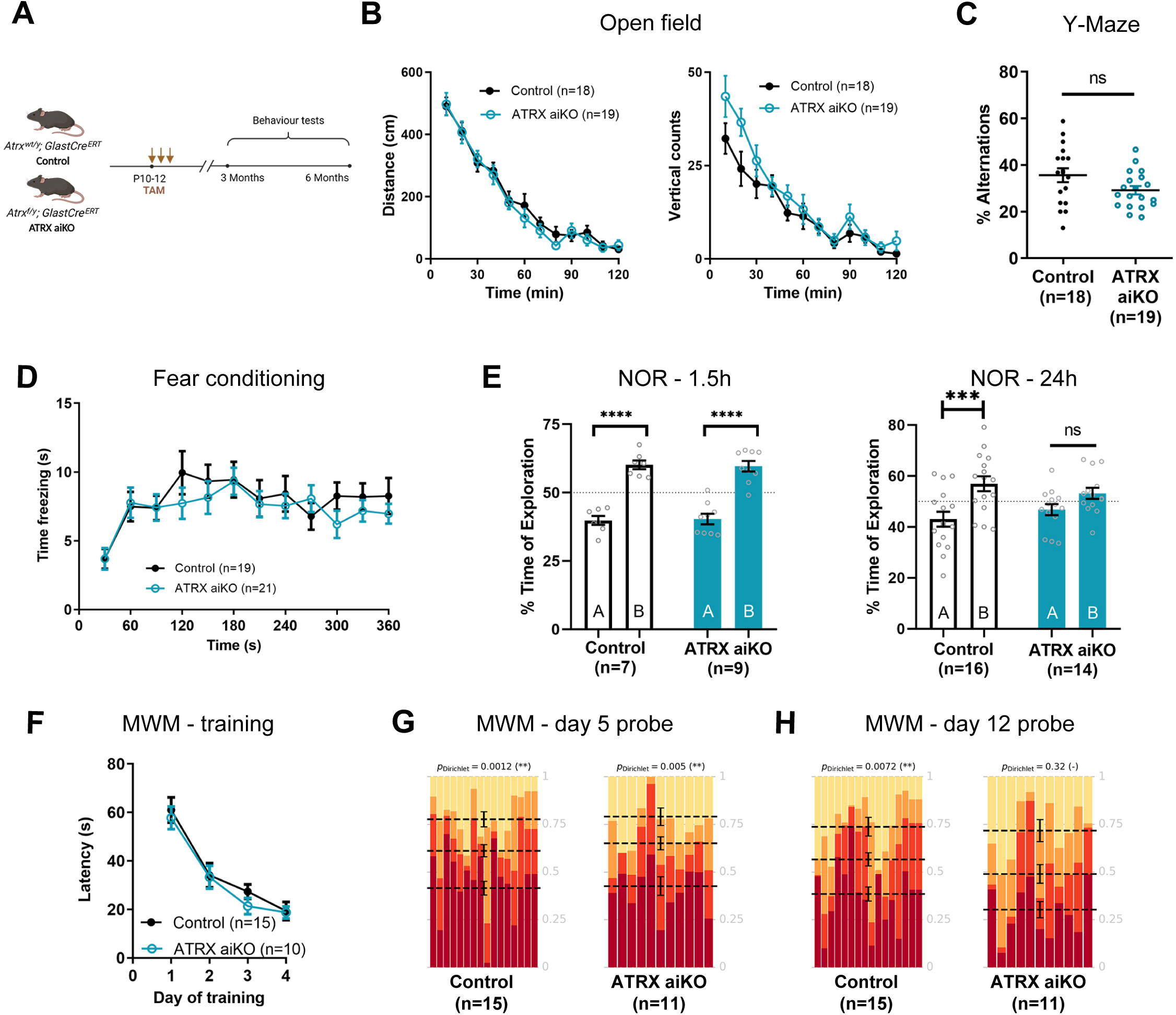
ATRX deficiency in astrocytes leads to long-term spatial and recognition memory deficits. (A) Experimental outline. (B) Left panel, distance travelled during 120 minutes in the open field test, twANOVA F_Genotype (1,35)_=0.237, p=0.629. Right panel, vertical episodes during 120 minutes in the open field test, twANOVA F_Genotype (1,35)_=2.307, p=0.138. (C) Alternations during the 5 min Y-maze test for working memory, unpaired Student’s t-test, t_(35)_=1.884, p=0.068). (D) Time freezing in the contextual fear conditioning paradigm, 24 h post-shock, twANOVA, F_Genotype (1,38)_=0.369, p=0.547. (E) Time spent exploring training object A and the novel object B in the novel object recognition (NOR) test 1.5 h and 24 h post-training. 1.5h: twANOVA, F_Novelty(1,28)_=113.3 p<0.0001, F_Genotype(1,28)_=8.92×10-14 p>0.9999; mc****p<0.0001), 24h: twANOVA, F_Novelty(1,56)_=14.71 p=0.0003, F_Genotype(1,56)_=7.306×10^-14^ p>0.999; mc***p<0.001). (F) Latency to find the hidden platform in the Morris water maze task over four training days. twANOVA, F(1, 30)=0.081, p=0.778. (G) Fraction of time spent in each quadrant during the probe day 5, Dirichlet uniformity test, Control = 15.8662, p=0.0012 and ATRX aiKO = 12.8263, p=0.0050. (H) Fraction of time spent in each quadrant during the probe day 12, Dirichlet uniformity test, Control = 13.6215, p=0.0035 and ATRX aiKO = 3.4849, p=0.3227. A-F data depicted as mean ± SEM. mc, multiple comparisons. G and H, means per quadrant shown with the doted lines, target quadrant shown at the bottom of the plots. Number of mice (n) shown in each experiment.

We next tested memory in the novel object recognition task. Both control and ATRX aiKO mice performed equally well during training, showing no baseline preference for either of the two identical objects (**Fig. S3E,H**). In the short-term probe test (1.5 h), ATRX aiKO mice spent significantly more time with the novel object, exhibiting similar discrimination index as control mice (**Fig. 5E and S3F**). Conversely, at 24 h, ATRX aiKO mice no longer showed a preference for the novel object, indicating a deficit in long-term recognition memory (**Fig. 5E**, **S3G**). This difference was not due to altered exploration time (**Fig. S3H**). Spatial learning and memory were assessed in the Morris water maze paradigm. There was no difference in the latency to complete the learning task over 4 days of training (**Fig. 5F**). During the probe test on the fifth day, control and ATRX aiKO mice spent significantly more time in the target quadrant, suggesting intact short-term spatial memory (**Fig. 5F**). However, after one week (probe day 12), ATRX aiKO mice did not spend more time in the target compared to other quadrants, suggesting long-term spatial memory deficits (**Fig. 5F**). Summary of all statistical analysis of the behavior assays is reported in **Table S3**. Collectively, behavioral assessments in ATRX aiKO mice identified specific deficiencies in long-term recognition and spatial memory processes traditionally linked to hippocampal function.

## Discussion

Astrocytes have emerged as central regulators of brain homeostasis and neuronal function, with increasing evidence linking their dysfunction to neurodevelopmental disorders. In this study, we demonstrate that selective loss of ATRX in astrocytes leads to decreased excitatory synaptic input and hyperexcitability in CA1 hippocampal neurons. These changes are accompanied by early, transient structural alterations in neurons that resolve by adulthood. Importantly, these cellular and physiological changes are associated with specific deficits in long-term spatial and recognition memory, underscoring the importance of chromatin regulation in astrocytes for cognitive processes.

Morphological analysis of ATRX-null astrocytes did not reveal significant alterations in the overall complexity of GFAP-positive astrocytes, although a modest increase in proximal branching was observed. The lack of pronounced morphological changes may be due to compensatory mechanisms by neighboring wild-type astrocytes or the limitations of GFAP-based tracing, which does not capture the full extent of astrocytic processes. Recent studies have emphasized the heterogeneity of astrocyte morphology both across and within brain regions (Endo 2022) and more comprehensive labelling and three-dimensional reconstruction approaches may be necessary to fully elucidate the structural consequences of ATRX loss.

We demonstrate that ATRX deletion in astrocytes led to dynamic, time-dependent changes in hippocampal gene expression that shifted from early cytoskeletal and immune alterations at one month to metabolic and ion transport pathway disruption by three months. This temporal progression aligns with emerging evidence that astrocytes undergo distinct maturation phases, with early postnatal periods representing windows of heightened vulnerability to chromatin disruption [20]. The early enrichment for immune response categories could reflect altered astrocyte-microglia crosstalk during development [21, 22]. Astrocytes are key regulators of cerebral metabolism, and disruptions in glycolytic pathways, redox balance, and mitochondrial function have been linked to neuronal dysfunction. Our findings suggest that ATRX deficiency perturbs these critical astrocytic functions, which may impact neuronal excitability and synaptic activity. Additionally, DEGs associated with potassium and glutamate handling implicate astrocytic regulation of the extracellular ionic environment and neurotransmitter clearance as potential mediators of the observed neuronal phenotypes.

Electrophysiological recordings showed that CA1 pyramidal neurons in the context of ATRX-deficient astrocytes exhibit increased intrinsic excitability and a reduced frequency of spontaneous excitatory postsynaptic currents. It is difficult to determine if the observed alterations in sESPC frequency appear in response to the increased action potential firing, but it is likely that these two phenomena are linked and point to a possible compensatory mechanism. Our results also show unaltered threshold and membrane resistance, indicating that neurons in the context of ATRX-null astrocytes possess normal overall number and conductance of ion channels. Astrocytes are regulators of extracellular ionic levels, and astrocyte-mediated disruption of this ionic balance is known to affect neuronal excitability [23]. Extracellular K^+^ is one of the mechanisms through which astrocytes can modulate neuronal excitability and contribute to disease (Reviewed in [24, 25]), and could be a mechanism by which ATRX-null astrocytes affect neuronal function in the ATRX aiKO hippocampus.

Behavioral analyses revealed that ATRX loss in astrocytes selectively impairs long-term hippocampal-dependent memory, while short-term memory remains unaffected. This specificity is in line with previous studies that implicate astrocytes in memory consolidation and retrieval (Reviewed in [26–28]). For example, transgenic mice designed to specifically impede astrocyte signaling exhibit altered cortical gamma oscillations and impaired recognition memory [29, 30]. Moreover, astrocyte-specific deletion of cannabinoid receptor 1 also leads to long-term recognition memory deficits in mice [31] in mice (Robin et al., 2018). Astrocytes have been shown to regulate long-term memory consolidation and retrieval by releasing molecules including adenosine [32], D-serine [33] and BDNF [34] as well as metabolic substrates like lactate [35] The preservation of working memory, contextual fear conditioning, and anxiety-related behaviors suggests that ATRX’s role in astrocytes is selectively required for specific memory processes rather than global cognitive or emotional function.

The moderate efficiency of ATRX deletion achieved in this study may underestimate the full spectrum of phenotypes associated with complete astrocytic loss. The ∼50% recombination efficiency using these Cre system may depend on when Cre translocation is induced, since the values reported by other groups using the same Cre driver line vary [36, 37] and likely reflects GLAST expression patterns at that time point and overall astrocyte heterogeneity [38–42] Yet, robust functional and behavioral phenotypes were observed, suggesting that astrocyte networks are more vulnerable to disruption than previously thought.

Our findings demonstrate that ATRX-mediated chromatin regulation in astrocytes is essential for maintaining hippocampal neuronal function and long-term memory. These results provide a mechanistic framework for understanding how astrocytic dysfunction may contribute to intellectual disability in ATR-X syndrome and highlight astrocytes as promising therapeutic targets for cognitive disorders. Further dissection of astrocyte-specific molecular pathways will be critical for developing targeted interventions in neurodevelopmental disease.

## Methods

### Mouse husbandry

All procedures involving animals were conducted in accordance with the regulations of the Animals for Research Act of the province of Ontario and approved by the University of Western Ontario Animal Care and Use Committee. Mice were exposed to 12-hour light/12-hour dark cycles and fed *ad libitum* with tap water and regular chow. The *Atrx*^loxP^ mice have been described previously [11] (MGI:3528480). To delete ATRX in astrocytes, C57BL/6 heterozygous *Atrx*^loxP^ females were mated with heterozygous B6 Glast-CreER [12] (Tg(Slc1a3-cre/ERT)1Nat, IMSR JAX:012586, MGI:4430111) male mice that express inducible Cre recombinase under the control of the *Slc1a3* (Glast) promoter. We also introduced the Cre-sensitive tdTomato-Ai14 allele (Ai14, B6;Cg-Gt(ROSA)26Sor^tm14(CAG-tdTomato)Hze^, IMSR JAX:007908 MGI:3809524) to fluorescently label Cre-positive astrocytes. To isolate fluorescent-tagged astrocytic nuclei, we incorporated the Cre-sensitive Sun1-GFP allele, obtained from the Jackson Laboratories (B6;129-Gt(ROSA)26Sor^tm5(CAG-Sun1/sfGFP)Nat/J^, IMSR JAX:021039, MGI:5614796) [43]. The Thy1-GFP-M allele (B6;Tg(Thy1-EGFP)MJrs, IMSR JAX:007788, MGI:3766828) was also introduced to sparsely label neurons. Genomic DNA from ear punches or tail biopsies was isolated for genotyping by PCR amplification using the primers listed in **Table S4**. PCR conditions were 3 min at 95 °C, 35 cycles at 95 °C for 10 s, 57 °C for 20 s, 72 °C for 60 s, and 5 min at 72 °C.

### Tamoxifen treatment

10 mg tamoxifen (Cat# T5648, Sigma) was dissolved in 100 µL 95% ethanol at 65°C and diluted in 900 µL corn oil (Cat# C8267, Sigma). Pups were injected intraperitoneally with 1 mg tamoxifen starting at post-natal day 10 for three consecutive days.

### Behavioral assessments

All behavioral tasks were performed between 9:00 AM and 7:00 PM. ARRIVE guidelines were followed. Mouse groups were randomized, experimenters were blind to the genotypes, and software-based analysis was used to score mouse performance in all the tasks. Mice were habituated in their home cages for 30 minutes prior to testing. All behavioral assays were performed when mice were between 3-7 months of age. Multiple cohorts of male mice (3-6) were tested independently to reach the final sample size, starting with less demanding tasks and ending with more demanding ones. Video recording and tracking were performed using AccuScan Instrument and AnyMaze software.

#### Open field

Mice were placed in a 20 cm x 20 cm arena with 30 cm high walls. Locomotor activity was measured in 5 min intervals over a 2 h period, as previously described [44].

#### Light-dark box test

Mice were placed in an arena with one half covered (dark compartment). Locomotor activity was measured for 5 minutes, and the time spent in each compartment was recorded.

#### Y-Maze test

Spontaneous alternation was measured using a three-armed Y-maze as previously described [45]. Video tracking was performed for 5 min and spontaneous alternations (entry in three different arms without visiting a previous arm) were recorded.

#### Novel object recognition test

Mice were habituated in a 40 cm x 40 cm arena with walls of 30 cm height for 5 min for two consecutive days. On day 3, mice were placed in the arena with two identical objects (A) and allowed to explore freely for 10 minutes. Animals were randomly assigned to one of two test probes, a short-term memory test at 1.5 h after training, or a long-term memory test at 24 h post-training. For probe tests, mice were placed in the arena with the training object (A) and a novel object (B) and allowed to explore for 5 minutes. The amount of time spent exploring each object was tracked manually from videos. Exploration time included time sniffing or touching the object, but not leaning or climbing on the object. Recognition memory index was calculated as the percentage of time spent with each object as a fraction of the total time exploring objects. Only animals displaying at least 10 s exploration were used in calculations. The objects used and location in the arena were counterbalanced as training or novel object between the groups.

#### Contextual fear memory

Mice were placed in a 10 cm x 20 cm clear acrylic enclosure with 30 cm high walls with drawings of black stripes and a black star on the opposite wall. The metal grid floor was equipped with an electric shock generator. On the first day, mice were placed in the enclosure and left to explore for 3 min, with a shock (2 mA, 180 V, 2 s) given at 2.5 min. The next day, mice were placed back in the same enclosure for 6 min. Freezing (lack of motion anywhere in the body and overall movement) and immobility (lack of movement in the arena) were measured.

#### Morris water maze

A 1.5 m diameter tank with 24 °C water and a platform submerged approximately 1.5 cm beneath the surface was used for the task. Shapes printed in black and white distributed around the tank were used as spatial cues. For training, mice performed four 90 s trials, with a 15 min intertrial period, daily at the same time of day for 4 consecutive days. If the mice did not find the platform within 90 s, they were gently guided onto the platform. On days 5 and 12, the platform was removed, and mice were placed in the tank for 1 min. Latency, distance, and speed to find the platform were recorded for training trials. Time spent in each quadrant was recorded for probe trials on days 5 and 12. All animals performed both probe trials. Data was analyzed using the Dirichlet distribution test, using a pipeline specifically designed for the Morris water maze task [46].

### Slice preparation and electrophysiology

Mice were anesthetized using sodium pentobarbital (100mg/kg intraperitoneally) and transcardially perfused with cold (2–4 °C) NMDG-HEPES solution containing (in mM): 92 NMDG, 93 HCl, 2.5 KCl, 1.2 NaH_2_PO_4_, 30, NaHCO_3_, 20 HEPES, 25 Glucose, 5 sodium ascorbate, 2 Thiourea, 3 sodium pyruvate, 10 MgCl_2_, 0.5 CaCl_2_ (300–310 mOsm), saturated with 95% O_2_/5%CO_2_. Brains were quickly removed and placed in cold NMDG-HEPES solution for slicing. Coronal sections (350 μm thick) containing the hippocampus were cut using a vibratome (VT-1200, Leica Biosystems). Slices were trimmed around CA1 and incubated at 34 °C for 15 minutes in NMDG-HEPES solution saturated with 95% O_2_/5%CO_2_. Slices were then transferred to artificial cerebrospinal fluid (aCSF) consisting of (in mM): 126 NaCl, 2.5 KCl, 26 NaHCO_3_, 2.5 CaCl_2_, 1.5 MgCl_2_, 1.25 NaH_2_PO_4_ and 10 D-glucose (295–300 mOsm), saturated with 95% O_2_/5%CO_2_ and maintained at room temperature. Slices were transferred to a recording chamber superfused with aCSF at a flow rate of 1.5–2.0 mL/min and maintained at 27–30 °C. CA1 neurons were visualized using an upright microscope with infrared differential interference contrast optics (BX 51WI, Olympus). Borosilicate glass recording pipettes (BF120-69-15, Sutter Instruments) were pulled in a Flaming/Brown Micropipette Puller (P-1000, Sutter Instruments) with a resistance between 3–5 MΩ. Pipettes were filled with an internal solution containing (in mM): 116 K-gluconate, 8 KCl, 12 Na-gluconate, 10 HEPES, 2 MgCl_2_, 4 K_2_ATP, 0.3 Na_3_GTP and 1 K_2_-EGTA (283–289 mOsm, pH 7.2–7.4). Glutamatergic sEPSCs were isolated by adding picrotoxin (100 μM) to the aCSF while holding the postsynaptic neuron at -80 mV in voltage clamp. Spike firing was measured in current clamp from a holding potential of -80mV using a step protocol from -160pA to 460pA in 40pA increments. Access resistance was monitored throughout the recording and cells were discarded if the value exceeded 20 MΩ. The reported membrane potential was corrected for a calculated junctional potential of -12mV.

### Patch clamp data collection, analysis and statistics

Whole cell patch clamp recordings were obtained using a Multiclamp 700B amplifier (Molecular Devices, California, USA), low pass filtered at 1 kHz and digitized at a sampling rate of 20 kHz using Digidata 1440A (Molecular Devices). Data was recorded on a PC using pClamp 10.6 (Molecular Devices) and analysed using MiniAnalysis (Synaptosoft, Georgia, USA) for EPSCs, Clampfit (Molecular Devices) for membrane potential and a custom Python algorithm (adapted from a white paper from Allen Cell Types Database) for cell firing. Briefly, the slope (dV/dt) was measured by taking the difference in voltage between two time steps and dividing by the resolution of acquisition. The time of the threshold crossing was detected by finding the time point where dV/dt was ≤5% of the maximum dV/dt of the rising phase. The following criteria were used to detect action potentials during current injection steps: 1) the duration from threshold to peak is ≤5 ms, 2) amplitude is ≥2mV and absolute peak ≥-30mV, and 3) action potential trough (minimum membrane potential in the interval between the peaks of two consecutive action potentials) is ≤–22 mV. Statistical tests were performed in GraphPad Prism 7 (GraphPad Software Inc, California, USA) and p < 0.05 was considered as statistically significant. Two group comparisons were done using paired t-tests. For sEPSC analysis, baseline data was taken at least 5 mins after breaking through into whole-cell mode and a 0.5- or 1-min bin was used for analysis. To achieve an accurate measure of the amplitude, individual sEPSCs were visually screened in MiniAnalysis and events below 5 pA were not included.

### CA1 neuron tracing and spine analysis

Mice were transcardially perfused with PBS followed with 4% paraformaldehyde (PFA) in PBS. Brains were fixed for 48h in PFA and sectioned at 150 µm thickness on a Vibratome Series 1000. Sections were mounted and imaged on a laser scanning confocal microscope (Leica TCS SP8 and Nikon Eclipse Ti2). Z-stacks of hippocampal CA1 pyramidal neurons were obtained (3-4 µm thickness, 20-30 z intervals). Dendrites were traced using the Simple Neurite Tracer ImageJ plugin (FIJI) and analyzed in a blinded manner using the Sholl plugin in FIJI at a radius step size of 4 µm as previously described [47]. Five to 10 neurons per animal were imaged, obtained from 6 pairs of 1-month-old and 5 pairs of 3-month-old mice of each genotype. For spine analysis, apical dendrites of hippocampal CA1 neurons were imaged with a Nikon Eclipse Ti2 confocal microscope with a 63X oil immersion objective. Z-stacks of 80 to 120 slices of 0.5 µm thickness were transformed to image sequences for analysis using Reconstruct software [48]. The length and width of spines from segments of secondary and tertiary dendrites were measured and used to classify spine morphology according to Risher et al. [4][4]. Five dendrite segments per neuron, from 5 to 10 neurons per mouse were analyzed, using 3 pairs of 3-month-old mice from each genotype.

### Immunofluorescence microscopy and imaging

Mice were transcardially perfused with PBS followed with 4% PFA in PBS. Brains were fixed overnight in PFA, washed with PBS the next day and cryopreserved by transferring to 30% sucrose in PBS at 4 °C until sunk. Brains were frozen on dry ice using Cryomatrix (Cat# 6769006, ThermoFisher) and sectioned in a cryostat (Leica CM 3050S) at 10 µm thickness on Superfrost plus slides (Cat# 22-037-246 ThermoFisher) and stored at -80 °C as described previously (Watson et al 2013). For immunofluorescence staining, slides were thawed and rehydrated in PBS, washed in PBS + 0.1% TritonX-100 (Cat# T8787 Millipore Sigma), blocked with 5% normal goat serum (Cat# G9023, Millipore Sigma), 0.5% BSA (Cat# ALB001.50 Bioshop) and 0.1% TritonX-100 in PBS for 1 h and incubated with primary antibodies overnight at 4 °C. Slides were washed three times with washing solution, incubated with secondary antibody for 1 h in blocking solution in the dark, washed twice, and counterstained with 1 µg/mL DAPI (Millipore Sigma D9542) for 5 min before mounting with Permafluor (Thermo Fisher TA-006-FM) or Immu-Mount (Thermo Fisher 9990402). The primary antibodies used are listed in **Table S5.** Slides were imaged using an inverted microscope (DMI 6000b, Leica) with digital camera (ORCA-ER, Hamamatsu). Images were captured with Openlab (PerkinElmer 5.0, RRID:SCR_012158) and processed using Volocity (PerkinElmer 6.0.1, RRID:SCR_002668) and Adobe Photoshop CS5 12.0. Cell counts in the cortex and hippocampus were performed in a blinded manner using the Cell count plugin in ImageJ. At least two sections per mouse from 3 pairs of mice of each genotype were used for counting.

### RNA extraction and library preparation

RNA was extracted from mouse hippocampus using the PureLink™ RNA Mini Kit (cat# 12183020, Thermo Fisher Scientific) with DNAse treatment (Cat# 12185010, Thermo Fisher Scientific). PolyA+ libraries were prepared from 1.5 µm of RNA at Canada’s Michael Smith Genome Sciences Centre (BC Cancer Research, Vancouver, BC, Canada, https://www.bcgsc.ca).

### Next-generation RNA sequencing and data analysis

Raw paired-end RNA-seq reads were trimmed using Trim Galore (v0.6.6) with the following parameters: --phred33 --length 36 -q 5 --stringency 1 -e 0.1. Trimmed reads were aligned to the *Mus musculus* GRCm38.p6 primary assembly (Ensembl release) using HISAT2 (v2.0.4) with default settings [49]. SAM files were sorted and converted to BAM format using SAMtools (v1.9) [50, 51]. Transcript assembly and quantification were performed for each sample with StringTie (v2.1.5), using the *Mus musculus* GRCm38 genome annotation as reference [52]. Transcript-level abundances were imported into R using the tximport (v1.20.0) R/Bioconductor package.

Differential expression analysis was performed at the transcript level using DESeq2 (v1.30.1)[53]. Lowly expressed transcripts (sum of counts ≤ 1 across samples) were filtered out prior to analysis. P values were weighted and adjusted for multiple testing using the Independent Hypothesis Weighting (IHW) method (adjusted p-value <= 0.05) [54]. Lancaster aggregation was then used to combine transcript-level p-values applied with a weighting scheme based on mean expression level for the transcript to derive gene-level significance[55].

### Gene Ontology Enrichment Analysis

Gene Ontology (GO) enrichment analysis was performed using the topGO R package (v2.44.0). For each gene set of interest (e.g., genes with aggregated Lancaster p-value < 0.05), genes were mapped to the Biological Process (BP) ontology using the org.Mm.eg.db R/Bioconductor annotation package. Enrichment was assessed using Fisher’s exact test with the classic and weight01 algorithms. The full set of expressed genes in the dataset was used as the background. For each significant GO term, associated gene lists were extracted, and the top enriched terms were reported.

### Statistical analysis and data representation

Data analysis was performed using GraphPad Prism 7 or 8 (GraphPad Software Inc.), using Student’s t test to compare pairs of datasets. To compare multiple groups, one-way or two-way ANOVA for one or multiple variables was applied, respectively. P less or equal to 0.05 was considered significant. All results are depicted as ± SEM and post-hoc tests applied are indicated in the figure legends. Schematics done in Biorender.com

## Abbreviations

aCSF: Artificial cerebrospinal fluid
Ai14: B6.Cg-Gt(ROSA)26Sortm14(CAG-tdTomato)Hze/J
ATP: Adenosine triphosphate
ATRX: Alpha-thalassemia/mental retardation, X-linked
BDNF: Brain-derived neurotrophic factor
BSA: Bovine Serum Albumin
CA1: Cornu ammonis 1
CNS: Central nervous system
CTCF: CCCTC-binding factor
Cx: Cortex
DAPI: 4′,6-diamidino-2-phenylindole
DEG: Differentially expressed gene
GFAP: Glial fibrillary acidic protein
GSEA: Gene Set Enrichment Analysis
Hc: Hippocampus
NMDG-HEPES: N-methyl-d-glutamine 4-(2-hydroxyethyl)-1-piperazineethanesulfonic acid
PAP: Perisynaptic astrocytic processes
PBS: Phosphate-buffered saline
PCR: Polymerase chain reaction
PFA: Paraformaldehyde
RNA: Ribonucleic acid
RT-qPCR: Real-time quantitative polymerase chain reaction
TF: Transcription factor

## Acknowledgements

We thank Matthew Cowan at the Neurobehaviour Core Facility at the Robarts Research Institute. We also thank Yan Jiang for technical support. This work was supported by operating funds from the Canadian Institutes for Health Research (CIHR) to NGB and WI (FRN# 178329). This work was supported by BrainsCAN through the Canada First Research Excellence Fund.

## Data availability

ATRX aiKO and control hippocampal sequencing data can be accessed at NCBI Bioproject PRJNA1002364. Source data are provided with this paper.

## Author contributions

M.A.PO. contributed to conceptualization, data collection, formal analysis, and writing, reviewing, and editing of the original draft. J.S. contributed to data collection, formal analysis, and writing, reviewing, and editing of the original draft. S.S., A.G. and M.R. contributed to data collection and formal analysis. H.M. contributed to conceptualization and data collection. V.D. contributed to conceptualization, data collection, formal analysis, and writing, reviewing, and editing of the original draft. W.I. contributed to conceptualization, funding acquisition, supervision, and writing, reviewing, and editing of the original draft. N.G.B. contributed to conceptualization, formal analysis, funding acquisition, supervision, and writing, reviewing and editing of the original draft.

## Figure legends

**Figure S1:**
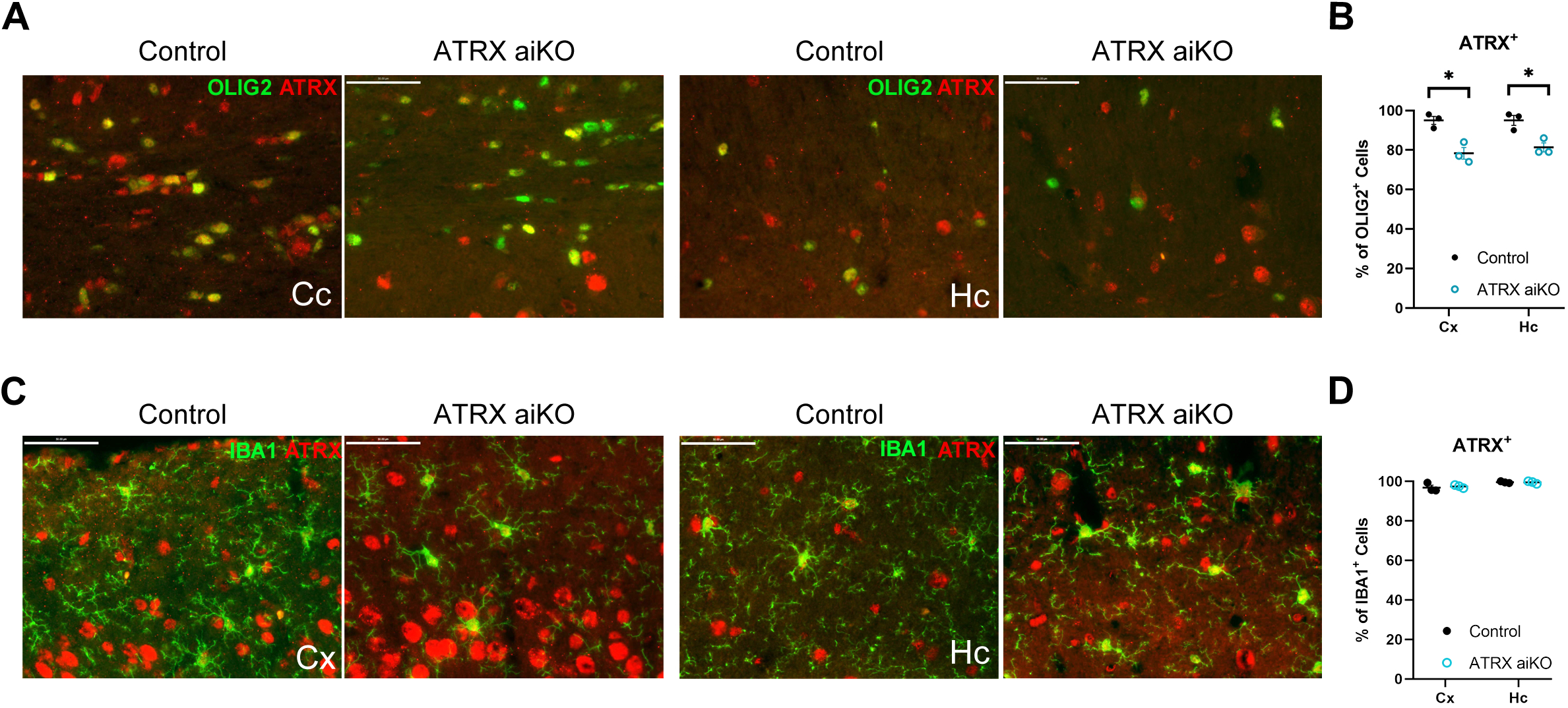
ATRX expression is largely retained in oligodendrocytes and microglia of ATRX aiKO mice. (A) Representative images of brain cryosections immunostained for ATRX and OLIG2 in the corpus callosum and hippocampus of control and ATRX aiKO mice. (B) Quantification of ATRX^+^OLIG2^+^ cells in the corpus callosum and hippocampus. (C) Immunofluorescence staining of ATRX and IBA1 in the cortex and hippocampus in control and ATRX aiKO brain cryosections. (D) Quantification of ATRX^+^IBA1^+^ cells. Scalebars, 100 µm. Cc: corpus callosum, Cx: cortex, Hc: hippocampus. Data shown as mean +/- SEM, n=3 control and ATRX aiKO brains. *p<0.05, unpaired Student’s t-test.

**Figure S2.**
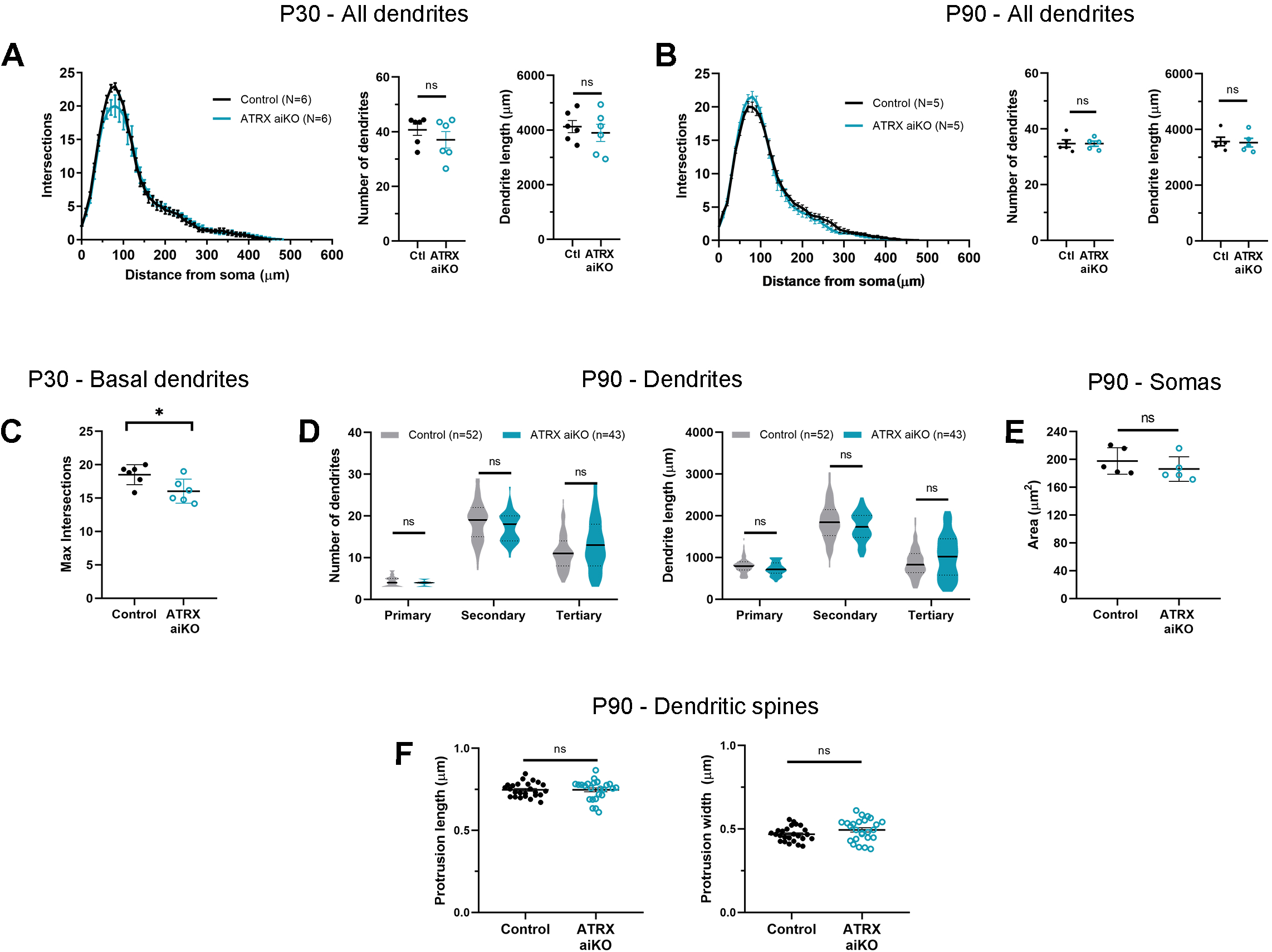
Dendrite and spine morphology in ATRX aiKO mice. (A) Sholl analysis of all CA1 dendrites at P30, twANOVA, F_Genotype (1,10)_=0.119, p=0.737. Total number of dendrites and total dendrite length, n=5-10 neurons from N=6 pairs of control and ATRX aiKO mice. (B) Sholl analysis of all CA1 dendrites at P90, twANOVA, F_Genotype (1,8)_=0.091, p=0.770. Total number of dendrites and total dendrite length, n=5-10 neurons from N=5 mice of each genotype. (C) Maximum number of intersections for CA1 basal dendrites at P30, n=5-10 neurons from N=6 mice of each genotype, unpaired Student’s t test, t_(10)_=2.582, p=0.0273. (D) Violin plots of number and length of primary, secondary and tertiary CA1 dendrites at P90. twANOVA F_Type(2, 279)_= 279.2, p<0.0001; F_Genotype(1, 279)_=0.00917, p=0.9237. n is the number of neurons analysed from N=5 mice of each genotype. (E) Soma area of hippocampal CA1 neurons at P90, n=10-30 neurons from N=6 mice of each genotype, unpaired Student’s t-test, t_(8)_=0.992, p=0.351. (F) Average spine length and width of apical CA1 dendrites at P90. Average of 5 segments from 5-10 neurons from N=3 mice of each genotype were analysed. Unpaired Student’s t-test was applied unless otherwise indicated. All data is depicted as +/- SEM.

**Figure S3:**
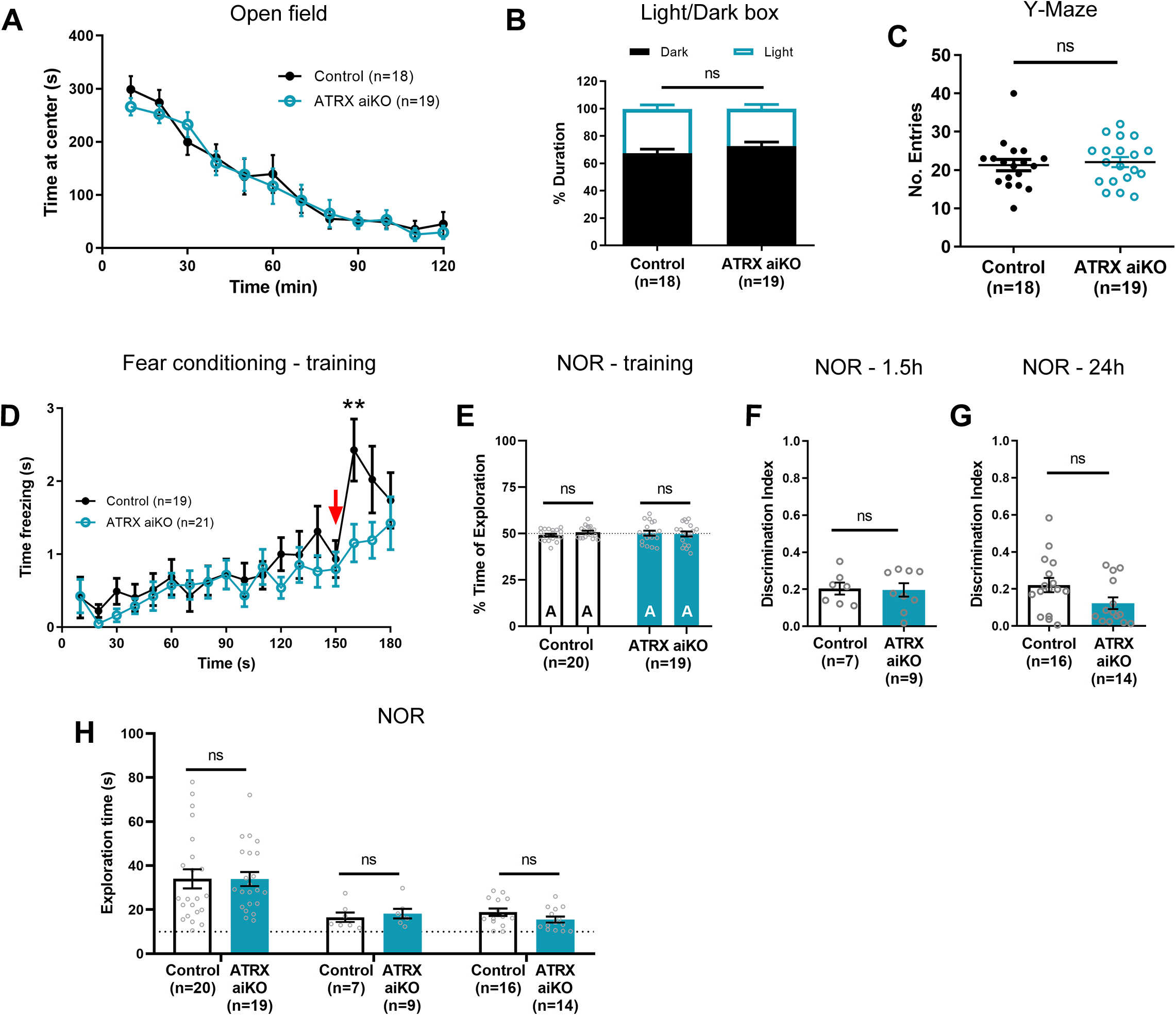
Astrocytic ATRX deficiency has no effect on anxiety, working or associative memory. (A) Time spent at the centre of the arena during 120 minutes in the open field test, twANOVA F_Genotype(1,35)_=0.066, p=0.798. (B) Time spent in the light and dark sections in the light/dark box test. twANOVA, F_Genotype(1,70)_=0.002, p_Dark_=0.969, p_Light_=0.452. (C) Number of arm entries during the 5 min Y-maze test, unpaired Student’s t-test, t_(35)_=0.391 p=0.698. (D) Time spent freezing during training in the contextual fear conditioning paradigm. Arrow indicates time of shock administration, twANOVA, F_Genotype(1,38)_=1.419, p=0.241; mc **p<0.01. (E) Percent of time spent exploring training object A in the novel object recognition task, twANOVA, F_Novelty(1,72)_=34.25 p<0.0001, F_Genotype(1,28)_=6.283×10^-14^, p>0.999. (F) Discrimination ratio for the 1.5 h probe, unpaired Student’s t-test, t_(14)_=0.138 p=0.892. (G) Discrimination ratio for the 24 h probe, unpaired Student’s t-test, t_(28)_=0.920 p=0.065. (H) Exploration time during novel object recognition training probe tests, unpaired Student’s t-test: Training, t_(46)_=1.046, p=0.301; 1.5 h probe, t_(20)_=0.196, p=0.847; 24 h probe, t_(34)_=1.024 p=0.313. All data is depicted as mean ± SEM.

## Notes

https://www.ncbi.nlm.nih.gov/bioproject/?term=PRJNA1002364

### Competing Interest Statement

The authors have declared no competing interest.

## References

1. Liu X, Ying J, Wang X, Zheng Q, Zhao T, Yoon S, et al. Astrocytes in Neural Circuits: Key Factors in Synaptic Regulation and Potential Targets for Neurodevelopmental Disorders. Front Mol Neurosci. 2021;14:729273.

2. Welle A, Kasakow C V, Jungmann AM, Gobbo D, Stopper L, Nordström K, et al. Epigenetic control of region-specific transcriptional programs in mouse cerebellar and cortical astrocytes. Glia. 2021;69:2160–2177.

3. Villarreal A, Vogel T. Different Flavors of Astrocytes: Revising the Origins of Astrocyte Diversity and Epigenetic Signatures to Understand Heterogeneity after Injury. Int J Mol Sci. 2021;22.

4. Gibbons RJ, Brueton L, Buckle VJ, Burn J, Clayton-Smith J, Davison BC, et al. Clinical and hematologic aspects of the X-linked alpha-thalassemia/mental retardation syndrome (ATR-X). Am J Med Genet. 1995;55:288–299.

5. Gibbons RJ, Wada T, Fisher CA, Malik N, Mitson MJ, Steensma DP, et al. Mutations in the chromatin-associated protein ATRX. Hum Mutat. 2008;29:796–802.

6. Drané P, Ouararhni K, Depaux A, Shuaib M, Hamiche A. The death-associated protein DAXX is a novel histone chaperone involved in the replication-independent deposition of H3.3. Genes Dev. 2010;24:1253–1265.

7. Levy MA, Kernohan KD, Jiang Y, Bérubé NG. ATRX promotes gene expression by facilitating transcriptional elongation through guanine-rich coding regions. Hum Mol Genet. 2014;24:1824–1835.

8. Nguyen DT, Voon HPJ, Xella B, Scott C, Clynes D, Babbs C, et al. The chromatin remodelling factor ATRX suppresses R-loops in transcribed telomeric repeats. EMBO Rep. 2017;18:914–928.

9. Truch J, Downes DJ, Scott C, Gür ER, Telenius JM, Repapi E, et al. The chromatin remodeller ATRX facilitates diverse nuclear processes, in a stochastic manner, in both heterochromatin and euchromatin. Nat Commun. 2022;13.

10. Sarma K, Cifuentes-Rojas C, Ergun A, Del Rosario A, Jeon Y, White F, et al. ATRX directs binding of PRC2 to Xist RNA and Polycomb targets. Cell. 2014;159:869–883.

11. Bérubé NG, Mangelsdorf M, Jagla M, Vanderluit J, Garrick D, Gibbons RJ, et al. The chromatin-remodeling protein ATRX is critical for neuronal survival during corticogenesis. Journal of Clinical Investigation. 2005;115:258–267.

12. Slezak M, Göritz C, Niemiec A, Frisén J, Chambon P, Metzger D, et al. Transgenic mice for conditional gene manipulation in astroglial cells. Glia. 2007;55:1565–1576.

13. Soto JS, Jami-Alahmadi Y, Chacon J, Moye SL, Diaz-Castro B, Wohlschlegel JA, et al. Astrocyte–neuron subproteomes and obsessive–compulsive disorder mechanisms. Nature. 2023. 27 April 2023. 10.1038/s41586-023-05927-7.

14. Reemst K, Noctor SC, Lucassen PJ, Hol EM. The indispensable roles of microglia and astrocytes during brain development. Front Hum Neurosci. 2016;10.

15. Bernardinelli Y, Randall J, Janett E, Nikonenko I, König S, Jones EV, et al. Activity-dependent structural plasticity of perisynaptic astrocytic domains promotes excitatory synapse stability. Current Biology. 2014;24:1679–1688.

16. Perez-Alvarez A, Navarrete M, Covelo A, Martin ED, Araque A. Structural and functional plasticity of astrocyte processes and dendritic spine interactions. Journal of Neuroscience. 2014;34:12738–12744.

17. Risher WC, Ustunkaya T, Alvarado JS, Eroglu C. Rapid golgi analysis method for efficient and unbiased classification of dendritic spines. PLoS One. 2014;9.

18. Mather DE, Gunsett FC, Allen OB, Kannenberg LW. Estimation of phenotypic selection differentials for predicting genetic responses to ratio-based selection. Genome. 1988;30:838–843.

19. Cao P, Maximov A, Südhof TC. Activity-dependent IGF-1 exocytosis is controlled by the Ca(2+)-sensor synaptotagmin-10. Cell. 2011;145:300–311.

20. Lattke M, Goldstone R, Ellis JK, Boeing S, Jurado-Arjona J, Marichal N, et al. Extensive transcriptional and chromatin changes underlie astrocyte maturation in vivo and in culture. Nat Commun. 2021;12.

21. Matejuk A, Ransohoff RM. Crosstalk Between Astrocytes and Microglia: An Overview. Front Immunol. 2020;11.

22. Vainchtein ID, Molofsky A V. Astrocytes and Microglia: In Sickness and in Health. Trends Neurosci. 2020;43:144–154.

23. Kofuji P, Newman EA. Potassium buffering in the central nervous system. Neuroscience. 2004;129:1043–1054.

24. Bellot-Saez A, Kékesi O, Morley JW, Buskila Y. Astrocytic modulation of neuronal excitability through K+ spatial buffering. Neurosci Biobehav Rev. 2017;77:87–97.

25. Nwaobi SE, Cuddapah VA, Patterson KC, Randolph AC, Olsen ML. The role of glial-specific Kir4.1 in normal and pathological states of the CNS. Acta Neuropathol. 2016;132.

26. Santello M, Toni N, Volterra A. Astrocyte function from information processing to cognition and cognitive impairment. Nat Neurosci. 2019;22:154–166.

27. Adamsky A, Goshen I. Astrocytes in Memory Function: Pioneering Findings and Future Directions. Neuroscience. 2018;370:14–26.

28. Kol A, Goshen I. The memory orchestra: the role of astrocytes and oligodendrocytes in parallel to neurons. Curr Opin Neurobiol. 2021;67:131–137.

29. Lee HS, Ghetti A, Pinto-Duarte A, Wang X, Dziewczapolski G, Galimi F, et al. Astrocytes contribute to gamma oscillations and recognition memory. Proc Natl Acad Sci U S A. 2014;111.

30. Sardinha VM, Guerra-Gomes S, Caetano I, Tavares G, Martins M, Reis JS, et al. Astrocytic signaling supports hippocampal–prefrontal theta synchronization and cognitive function. Glia. 2017;65:1944–1960.

31. Robin LM, Oliveira da Cruz JF, Langlais VC, Martin-Fernandez M, Metna-Laurent M, Busquets-Garcia A, et al. Astroglial CB1 Receptors Determine Synaptic D-Serine Availability to Enable Recognition Memory. Neuron. 2018;98:935–944.e5.

32. Martin-Fernandez M, Jamison S, Robin LM, Zhao Z, Martin ED, Aguilar J, et al. Synapse-specific astrocyte gating of amygdala-related behavior. Nat Neurosci. 2017;20:1540–1548.

33. Adamsky A, Kol A, Kreisel T, Doron A, Ozeri-Engelhard N, Melcer T, et al. Astrocytic Activation Generates De Novo Neuronal Potentiation and Memory Enhancement. Cell. 2018;174:59–71.e14.

34. Liu JH, Zhang M, Wang Q, Wu DY, Jie W, Hu NY, et al. Distinct roles of astroglia and neurons in synaptic plasticity and memory. Mol Psychiatry. 2022;27:873–885.

35. Suzuki A, Stern SA, Bozdagi O, Huntley GW, Walker RH, Magistretti PJ, et al. Astrocyte-neuron lactate transport is required for long-term memory formation. Cell. 2011;144:810–823.

36. Higashimori H, Schin CS, Chiang MSR, Morel L, Shoneye TA, Nelson DL, et al. Selective Deletion of Astroglial FMRP Dysregulates Glutamate Transporter GLT1 and Contributes to Fragile X Syndrome Phenotypes In Vivo. Journal of Neuroscience. 2016;36:7079–7094.

37. Cheli VT, Santiago González DA, Wan Q, Denaroso G, Wan R, Rosenblum SL, et al. H-ferritin expression in astrocytes is necessary for proper oligodendrocyte development and myelination. Glia. 2021;69:2981–2998.

38. Chai H, Diaz-Castro B, Shigetomi E, Monte E, Octeau JC, Yu X, et al. Neural Circuit-Specialized Astrocytes: Transcriptomic, Proteomic, Morphological, and Functional Evidence. Neuron. 2017;95:531–549.e9.

39. Bayraktar OA, Bartels T, Holmqvist S, Kleshchevnikov V, Martirosyan A, Polioudakis D, et al. Astrocyte layers in the mammalian cerebral cortex revealed by a single-cell in situ transcriptomic map. Nat Neurosci. 2020;23:500–509.

40. Batiuk MY, Martirosyan A, Wahis J, de Vin F, Marneffe C, Kusserow C, et al. Identification of region-specific astrocyte subtypes at single cell resolution. Nat Commun. 2020;11.

41. Herrero-Navarro Á, Puche-Aroca L, Moreno-Juan V, Sempere-Ferràndez A, Espinosa A, Susín R, et al. Astrocytes and neurons share region-specific transcriptional signatures that confer regional identity to neuronal reprogramming. Sci Adv. 2021;7.

42. Lozzi B, Huang T-W, Sardar D, Huang AY-S, Deneen B. Regionally Distinct Astrocytes Display Unique Transcription Factor Profiles in the Adult Brain. Front Neurosci. 2020;14:61.

43. Mo A, Mukamel EA, Davis FP, Luo C, Henry GL, Picard S, et al. Epigenomic Signatures of Neuronal Diversity in the Mammalian Brain. Neuron. 2015;86:1369–1384.

44. Tamming RJ, Siu JR, Jiang Y, Prado MAM, Beier F, Bérubé NG. Mosaic expression of *Atrx* in the mouse central nervous system causes memory deficits. Dis Model Mech. 2017;10:119–126.

45. De Castro BM, Pereira GS, Magalhães V, Rossato JI, De Jaeger X, Martins-Silva C, et al. Reduced expression of the vesicular acetylcholine transporter causes learning deficits in mice. Genes Brain Behav. 2009;8:23–35.

46. Maugard M, Doux C, Bonvento G. A new statistical method to analyze Morris Water Maze data using Dirichlet distribution. F1000Res. 2019;8:1601.

47. Tamming RJ, Dumeaux V, Jiang Y, Shafiq S, Langlois L, Ellegood J, et al. Atrx Deletion in Neurons Leads to Sexually Dimorphic Dysregulation of miR-137 and Spatial Learning and Memory Deficits. Cell Rep. 2020;31.

48. Fiala JC. Reconstruct: A free editor for serial section microscopy. J Microsc. 2005;218:52–61.

49. Kim D, Paggi JM, Park C, Bennett C, Salzberg SL. Graph-based genome alignment and genotyping with HISAT2 and HISAT-genotype. Nat Biotechnol. 2019;37:907–915.

50. Li H, Handsaker B, Wysoker A, Fennell T, Ruan J, Homer N, et al. The Sequence Alignment/Map format and SAMtools. Bioinformatics. 2009;25:2078–2079.

51. Danecek P, Bonfield JK, Liddle J, Marshall J, Ohan V, Pollard MO, et al. Twelve years of SAMtools and BCFtools. Gigascience. 2021;10.

52. Kovaka S, Zimin A V, Pertea GM, Razaghi R, Salzberg SL, Pertea M. Transcriptome assembly from long-read RNA-seq alignments with StringTie2. Genome Biol. 2019;20:278.

53. Love MI, Huber W, Anders S. Moderated estimation of fold change and dispersion for RNA-seq data with DESeq2. Genome Biol. 2014;15:550.

54. Ignatiadis N, Klaus B, Zaugg JB, Huber W. Data-driven hypothesis weighting increases detection power in genome-scale multiple testing. Nat Methods. 2016;13:577–580.

55. Yi L, Pimentel H, Bray NL, Pachter L. Gene-level differential analysis at transcript-level resolution. Genome Biol. 2018;19:53.

