## Supplemental Files for "Astrocytic Chromatin Remodeler ATRX Gates Hippocampal Memory Consolidation through Metabolic and Synaptic Regulation"

*Supplementary data*

**Table S1: Gene Ontology analysis of P30 hippocampal RNA-seq**

| GO Biological Process category | Genes | log2FoldChange | Lancaster p |
|---|---|---|---|
| negative regulation of microtubule depolymerization | Dysf | 0.2936 | 2.04E-03 |
| | Gas2l1 | 0.1180 | 1.36E-04 |
| | Nav3 | -0.0252 | 7.33E-03 |
| | Mid1ip1 | -0.0353 | 4.11E-02 |
| | Specc1l | -0.0411 | 2.08E-03 |
| | Tpx2 | -0.0467 | 3.77E-02 |
| | Atxn7 | -0.2756 | 7.65E-04 |
| microtubule bundle formation | Map1s | 0.1519 | 7.31E-03 |
| | Gas2l1 | 0.1180 | 1.36E-04 |
| | Kif20a | 0.0884 | 1.20E-02 |
| | Clip1 | -0.0147 | 3.54E-04 |
| | Ttll6 | -0.2675 | 1.75E-02 |
| | Dnah8 | -0.8999 | 1.90E-02 |
| | Rp1l1 | -5.2600 | 1.65E-02 |
| positive regulation of toll-like receptor 3 signalling pathway | Cav1 | -0.0611 | 5.34E-03 |
| | Peli1 | -0.2014 | 2.99E-02 |
| | Wdfy1 | -0.5318 | 8.92E-04 |
| positive regulation of NF-kappaB transcription factor activity | Ripk4 | 1.1309 | 7.42E-05 |
| | Ripk2 | 0.8958 | 3.25E-13 |
| | Eda2r | 0.8443 | 3.93E-02 |
| | Nts | 0.5136 | 1.30E-02 |
| | Nlrp3 | 0.3963 | 3.64E-02 |
| | Cd36 | 0.1456 | 2.01E-02 |
| | Map3k13 | 0.1192 | 3.29E-02 |
| | Prkd2 | 0.0682 | 6.89E-03 |
| | Il18 | 0.0523 | 4.47E-02 |
| | Cav1 | -0.0611 | 5.34E-03 |
| | Rab7b | -0.5112 | 2.84E-02 |

| GO Biological Process category | Genes | log2FoldChange | Lancaster p |
|---|---|---|---|
| positive regulation of synapse assembly | Oxt | 7.8757 | 4.26E-03 |
| | Ube2v2 | 0.3288 | 2.53E-04 |
| | Lrtm2 | 0.1918 | 3.17E-02 |
| | Sema4a | 0.1214 | 1.85E-02 |
| | Bhlhb9 | -0.0316 | 2.37E-03 |
| | Amigo1 | -0.0492 | 1.19E-02 |
| | Cux2 | -0.0802 | 3.16E-03 |
| | Srpx2 | -0.1659 | 4.66E-02 |
| positive regulation of neuron migration | Tbc1d24 | 0.0402 | 3.87E-03 |
| | Pax6 | 0.0104 | 3.12E-04 |
| | Sema6a | -0.2401 | 4.35E-02 |
| | Reln | -0.3842 | 4.30E-02 |
| positive regulation of dendritic spine morphogenesis | Camk2b | 0.0569 | 2.10E-02 |
| | Bhlhb9 | -0.0316 | 2.37E-03 |
| | Cux2 | -0.0802 | 3.16E-03 |
| | Reln | -0.3842 | 4.30E-02 |

**Table S2: Gene Ontology analysis of P30 hippocampal RNA-seq**

| GO Biological Process category | Genes | log2FoldChange | Lancaster p |
|---|---|---|---|
| cellular response to glucose starvation | Brsk2 | 0.058395 | 0.000214 |
| | Chka | -0.13838 | 0.033575 |
| | Gck | 0.633199 | 6.03E-05 |
| | Kat5 | 0.046257 | 0.002606 |
| | Mtmr3 | 0.035175 | 0.010956 |
| | Nuak2 | -0.10069 | 0.005266 |
| | Plin3 | 0.608271 | 5.01E-14 |
| | Prkaa1 | -0.16012 | 0.016559 |
| | Sh3glb1 | -0.1245 | 0.005301 |
| | Sirt1 | -0.083 | 2.22E-06 |
| | Suv39h1 | -0.10618 | 0.015402 |
| | Trp53 | 0.001472 | 0.012945 |
| fructose 6-phosphate metabolic process | Fbp1 | 1.152856 | 0.00323 |
| | Fbp2 | 1.183366 | 0.012984 |
| | Gck | 0.633199 | 6.03E-05 |
| | Gnpda1 | -0.03623 | 2.55E-07 |
| | Hk2 | 0.235956 | 0.009324 |
| | Pfkl | 0.016928 | 0.000682 |
| fructose metabolic process | Aldoa | 0.04828 | 1.71E-09 |
| | Aldob | -0.45802 | 0.040618 |
| | Fbp1 | 1.152856 | 0.00323 |
| | Fbp2 | 1.183366 | 0.012984 |
| | Gnpda1 | -0.03623 | 2.55E-07 |
| | Pfkfb1 | 0.0375 | 0.045491 |

**Table S3: Statistical analysis of behaviour assays**

| Fig. | Groups | N | Measurements | Analysis | F | p | Post-hoc |
|---|---|---|---|---|---|---|---|
| 5B | Control<br>ATRX aiKO<br>/Time | 18<br>19 | Total distance<br>(Open field) | 2way-<br>ANOVA | Interaction F (11, 385) = 0.3752<br>Time F (11, 385) = 96.55<br>Genotype F (1, 35) = 0.2374 | P=0.9652<br>P<0.0001<br>P=0.6291 | Sidak's m.c.<br>Control – ATRX aiKO through time<br>None significant. |
| | Control<br>ATRX aiKO<br>/Time | 18<br>19 | Vertical counts<br>(Open field) | 2way-<br>ANOVA | Interaction F (11, 385) = 1.458<br>Time F (4.639, 162.4) = 40.15<br>Genotype F (1, 35) = 2.307 | P=0.1451<br>P<0.0001<br>P=0.1377 | Sidak's m.c.<br>Control – ATRX aiKO through time<br>None significant. |
| 5C | Control<br>ATRX aiKO | 18<br>19 | % Alternations<br>(Y-Maze) | Unpaired<br>t-test | t=1.884, df=35 | P=0.0679 | - |
| 5D | Control<br>ATRX aiKO<br>/Time | 19<br>21 | Time Freezing<br>(Contextual Fear<br>Conditioning) | 2way-<br>ANOVA | Interaction F (11, 418) = 0.7908<br>Time F (11, 418) = 6.129<br>Genotype F (1, 38) = 0.3689 | P=0.6494<br>P<0.0001<br>P=0.5472 | Sidak's m.c.<br>Control – ATRX aiKO through time<br>None significant. |
| 5E | Control<br>ATRX aiKO | 7<br>9 | % Time of exploration<br>(Novel Object<br>Recognition)<br>1.5h Probe | 2way-<br>ANOVA | Interaction F (1, 28) = 0.08122<br>Novelty F (1, 28) = 113.3<br>Genotype F (1, 28) = 8.922e-14 | P=0.7778<br>P<0.0001<br>P>0.9999 | Sidak's m.c.<br>A – B Control, P<0.0001<br>A – B ATRX aiKO, P<0.0001 |
| | Control<br>ATRX aiKO | 16<br>14 | % Time of exploration<br>(Novel Object<br>Recognition)<br>24h Probe | 2way-<br>ANOVA | Interaction F (1, 56) = 2.044<br>Novelty F (1, 56) = 14.71<br>Genotype F (1, 56) = 7.306e-14 | P=0.1584<br>P=0.0003<br>P>0.9999 | Sidak's m.c.<br>A – B Control, P=0.0006<br>A – B ATRX aiKO, P=0.1991 |
| 5F | Control<br>ATRX aiKO | 15<br>10 | Latency<br>(Morris Water Maze)<br>Training | 2way-<br>ANOVA | Interaction F (3, 72) = 0.2239<br>Time F (2.294, 55.05) = 44.65<br>Genotype F (1, 24) = 0.4522 | P=0.8795<br>P<0.0001<br>P=0.5077 | Sidak's m.c.<br>Control – ATRX aiKO through time<br>None significant. |
| | Control<br>ATRX aiKO | 15<br>10 | Time in quadrant<br>(Morris Water Maze)<br>Probe Day 5 | Dirichlet<br>uniformity<br>test | Likelihood-ratio statistic (with<br>MWM correction)<br>Control = 15.8662<br>ATRX aiKO = 12.8263 | <br><br>P= 0.0012<br>P= 0.0050 | - |
| | Control<br>ATRX aiKO | 15<br>10 | Time in quadrant<br>(Morris Water Maze)<br>Probe Day 12 | Dirichlet<br>uniformity<br>test | Likelihood-ratio statistic (with<br>MWM correction)<br>Control = 13.6215<br>ATRX aiKO = 3.4849 | <br><br>P= 0.0035<br>P= 0.3227 | - |

**Table S4: List of primers**

| Gene | Use | Forward Primer | Reverse Primer |
|---|---|---|---|
| ATRX-wt | Genotyping | AGA AAT TGA GGA TGC TTC ACC | TGA ACC TGG GGA CTT CTT TG |
| ATRX-floxed | Genotyping | AGA AAT TGA GGA TGC TTC ACC | CCA CCA TGA TAT TCG GCA AG |
| Glast-CreER | Genotyping | ACA ATC TGG CCT GCT ACC AAA GC | CCA GTG AAA CAG CAT TGC TGT C |
| Sun1-GFP | Genotyping | AAG GGA GCT GCA GTG GAG TA | CGG GCC ATT TAC CGT AAG TTA T |
| TdTomato (Ai14) | Genotyping | GGC ATT AAA GCA GCG TAT CC | CTG TTC CTG TAC GGC ATG G |
| Thy1-EYFP | Genotyping | ACA GAC ACA CAC CCA GGA CA | CGG TGG TGC AGA TGA ACT T |

**Table S5: List of antibodies**

| Antibody | Species (Dilution) | Manufacturer | Cat#/ RRID |
|---|---|---|---|
| anti-ATRX | Rabbit polyclonal (1:100) | Santa Cruz Biotechnology | sc-15408 RRID:AB_2061023 |
| anti-ATRX | Mouse monoclonal (1:100) | Santa Cruz Biotechnology | sc-55584 RRID:AB_831012 |
| anti-GFAP | Mouse monoclonal (1:600) | Sigma-Aldrich Cat# | G3893 RRID:AB_477010 |
| anti-S100β | Rabbit polyclonal (1:200) | Agilent | Z0311 RRID:AB_10013383 |
| anti-Olig2 | Rabbit polyclonal (1:200) | Millipore | AB9610 RRID:AB_570666 |
| anti-Iba1 | Rabbit polyclonal (1:600) | Fujifilm Wako | 019-19741 RRID:AB_839504 |
